# Turbulence and blood washout in presence of mitral regurgitation: a computational fluid-dynamics study in the complete left heart

**DOI:** 10.1101/2023.03.19.533094

**Authors:** Lorenzo Bennati, Vincenzo Giambruno, Francesca Renzi, Venanzio Di Nicola, Caterina Maffeis, Giovanni Puppini, Giovanni Battista Luciani, Christian Vergara

**Affiliations:** Department of Surgery, Dentistry, Pediatrics, and Obstetrics/Gynecology, University of Verona, Verona, 37134, Italy; Division of Cardiac Surgery, Department of Surgery, Dentistry, Pediatrics, and Obstetrics/Gynecology, University of Verona, Verona, 37126, Italy; Department of Radiology, University of Verona, Verona, 37126, Italy; LaBS, Dipartimento di Chimica, Materiali e Ingegneria Chimica Giulio Natta, Politecnico di Milano, Milano, 20133, Italy

**Keywords:** turbulence, hemolysis, blood washout, mitral regurgitation, computational fluid dynamics, cine-MRI

## Abstract

In this work we performed a computational image-based study of blood dynamics in the whole left heart, both in a healthy subject and in a patient with mitral valve regurgitation (MVR). We elaborated dynamic cine-MRI images with the aim of reconstructing the geometry and the corresponding motion of left ventricle, left atrium, mitral and aortic valves, and aortic root of the subjects. This allowed us to prescribe such motion to computational blood dynamics simulations where, for the first time, the whole left heart motion of the subject is considered, allowing us to obtain reliable subject-specific information.

The final aim is to investigate and compare between the subjects the occurrence of turbulence and the risk of hemolysis and of thrombi formation. In particular, we modeled blood with the Navier-Stokes equations in the Arbitrary Lagrangian-Eulerian framework, with a Large Eddy Simulation model to describe the transition to turbulence and a resistive method to manage the valve dynamics, and we used a Finite Elements discretization implemented in an in-house code for the numerical solution.

Our results highlighted that the regurgitant jet in the MVR case gave rise to a large amount of transition to turbulence in the left atrium resulting in a higher risk of formation of hemolysis. Moreover, MVR promoted a more complete washout of stagnant flows in the left atrium during the systolic phase and in the left ventricle apex during diastole.

**NEW & NOTEWORTHY:** Reconstruction from cine-MRI images of geometries and motion of the left heart (left atrium and ventricle, aortic root, aortic and mitral valve) of a healthy and mitral regurgitant patient.

Prescription of such motion to a complete subject-specific computational fluid-dynamic simulation of the left heart. Investigation of turbulence in a regurgitant scenario.

Study of the mechanisms of prevention from stagnant flows and hemolysis formation in the atrium.

## Introduction

The pathologies affecting the left heart (LH) are the most common cause of death in the world [1]. One of these is mitral valve regurgitation (MVR), a condition leading to a formation of a regurgitant jet in the left atrium during the systolic phase due to an incomplete closure of the mitral valve leaflets. The formation and the development of the regurgitant jet may give rise to: i) presence of highly disturbed or even turbulent atrial flow that can lead to hemolysis in the atrium [2, 3] and ii) washing out of stagnant blood in the atrium that could prevent thrombi formation [4].

These phenomena are difficult to describe and quantify in the clinical practice. On the one hand, although clinical measures such as the regurgitant volume and the regurgitant fraction may provide significant information about the global cardiac function, they are not able to capture local features such as 3D velocity distribution and wall shear stresses [5, 6]. On the other hand, the space and time resolution of the available imaging techniques, such as Four-Dimensional Flow Magnetic Resonance Imaging (MRI) or Phase-Contrast MRI, is not nowadays enough accurate to capture small-scales features as recirculation areas, regions of transition to turbulence, and small coherent structures [7].

In this respect, computational methods can non-invasively provide quantitative information about the local pressure gradients, the velocity patterns and the shear forces, contributing to a better understanding of the cardiovascular system [8–14]. In particular, computational models applied to LH have contributed to a better knowledge of the LH physio-pathology [15–21] and to model and predict the outcomes of valve prostheses or surgical interventions [22–24]. Such methods can be broadly grouped in two categories: Fluid Structure Interaction (FSI) models and Dynamic Image-based Computational Fluid Dynamics (DIB-CFD) models with prescribed wall and valves motion. The latter approach has become, in the last decade, a valid alternative to FSI models when sufficiently detailed dynamic medical images are available [25–32]. In particular, regarding MVR, we mention works where the authors tested and compared different types of mitral valve prolapse [33], different degrees of MVR [34] and different functional changes of the ventricle and atrium in response to MVR [35]. However, no one of these studies was performed on a fully patient-specific LH geometry and displacement (ventricle + atrium + mitral valve + aortic valve + aortic root). Moreover, none of them investigated the transition to turbulence, nor the risk of hemolysis and the prevention from thrombi formation in the atrium.

The present study has four principal aims, which are all, at the best of our knowledge, innovative:

i. present a tool to perform a fully LH patient-specific DIB-CFD simulation with imposed motion of ventricle, atrium, aortic root, mitral valve, and aortic valve, acquired from cine-MRI images during the whole heartbeat;
ii. apply the previous tool in a healthy and in a MVR patient;
iii. describe and quantify the transition to turbulence in the MVR case;
iv. study the risk of hemolysis in the atrium and the prevention from thrombi formation with respect to the healthy subject. The significance of our results has been supported by the comparison with Echo Colour Doppler (ECD) measures in the healthy subject and with a cine-MRI flow pattern in the MVR case.

## Methods

In this section, we first described the cine-MRI images at disposal and the acquisitions of the ECD measurements; after, we detailed the reconstruction techniques used to obtain the patient-specific geometries and displacements of the left ventricle (LV), left atrium (LA), aortic root (AR), mitral valve (MV) and aortic valve (AV); then, we briefly reported the mathematical and numerical methods used in this work; finally we introduced the quantities of interest that has been analyzed in the Results section.

### Available cine-MRI images and ECD acquisitions

Cardiac cine-MRI data of two patients were provided by the Department of Radiology of Borgo Trento Hospital, Verona, Italy. Ethical review board approval and informed consent were obtained from all patients. In particular, we acquired dynamic images, consisting of 30 acquisitions per heartbeat, of a healthy subject (H) and of a patient with a severe MVR (R) due to a posterior leaflet prolapse. In Table 1, we reported some information about the two patients, including the heart rate, height, weight and body surface area (BSA) [36].

**Table 1.** For each patient, we reported the values of heart rate, height, weight and BSA.

| Patient | Heart rate [BPM] | Height [m] | Weight [kg] | BSA [$m^2$] |
| --- | --- | --- | --- | --- |
| H | 66 | 1.93 | 84 | 2.14 |
| R | 75 | 1.84 | 105 | 2.28 |

The acquisitions were performed with the Achieva 1.5T (TX) - DS (Philips, Amsterdam, Netherlands) technology. Specifically, for each patient, we have at disposal different acquisitions: the Short Axis series of LV, the Long Axis series and the Rotated series of MV (see [35] for further details). Moreover, for subject H we have at disposal also the Short Axis series of AR, consisting in volumetric series made of 4 slices with thickness and distancing of 8 *mm* along the aortic root main axis, with a spatial resolution of 1 *mm* and time resolution equal to 30 frames/cardiac cycle; see Figure 1, left. After the image acquisitions, patient R underwent to a surgical operation to restore a correct heart function.

**Figure 1.**
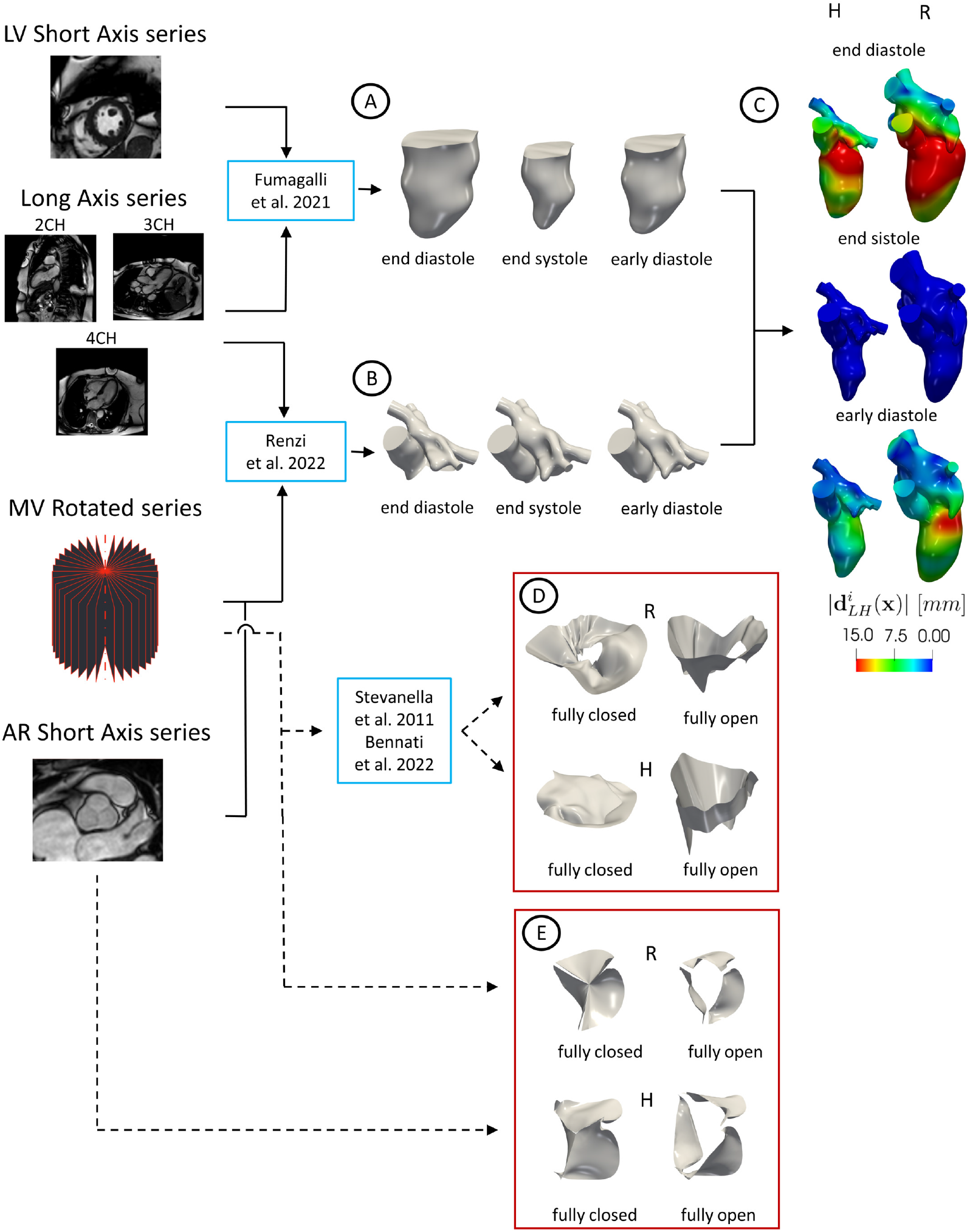
Flow chart to reconstruct the geometries of LH and valves and their displacements. Continuous lines refer to the steps followed to reconstruct the LH walls and the dashed lines to the reconstruction of the valves. A: geometric reconstruction of the LV endocardium in all the 30 frames by adopting the strategy of [32] (here we reported the geometries at three representative frames as an example). B: geometric reconstruction of AR and LA in all the 30 frames by adopting the strategy of [37] (here we reported the geometries at three representative frames as an example). C: merging of LV with AR and LA and registration of the displacement field 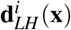 with respect to the end systolic configuration (here we reported the magnitude of 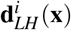 at three representative frames as an example). D: geometric reconstruction of MV in the fully open and closed configurations by adopting the method of [38]. E: geometric reconstruction of AV in the fully open and closed configurations.

We point out that the Long-Axis series and the Short-Axis series of LV are standard cine-MRI data that are routinely acquired in the clinical procedure. Instead, the Short-Axis series of AR and the Rotated series of MV represent ad hoc advanced cine-MRI acquisitions.

After the cine-MRI acquisitions, subject H underwent to ECD measurements with a EPIQ CVx ultrasounds scanner and linear 8MHz probe (Philips Ultrasound, Bothell, WA). The velocity measures were acquired at the peak instants at three different locations: *P*1: 1 *cm* upstream the AV base; *P*2: 1.6 *cm* downstream the MV annulus; *P*3: 5.5 *cm* far from the the LV apex, see Figure 2A.

**Figure 2.**
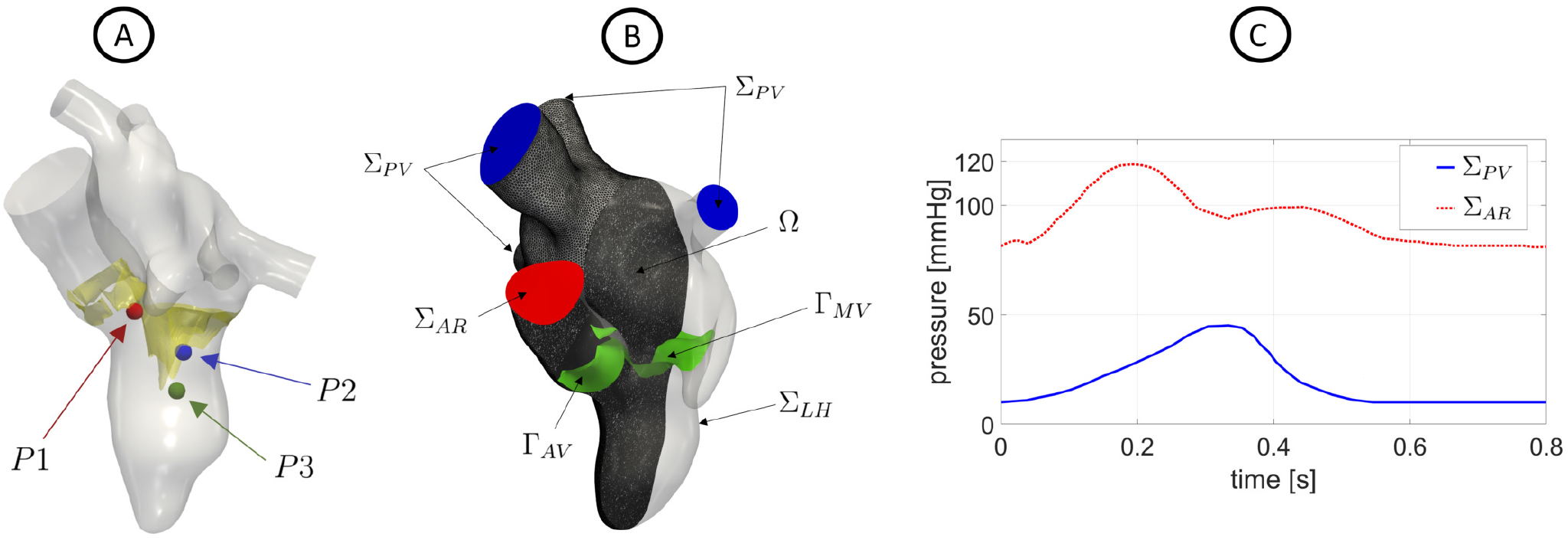
A: Location of the points where ECD measures were acquired. B: Computational domain Ω with its boundaries. In green we reported the aortic and mitral valves Γ_*AV*_ and Γ_*MV*_. The computational mesh of patient R is also displayed. C: trend in time of the pressures imposed at Σ_*PV*_ (for scenario R) and Σ_*AR*_ (for H, the curve at the AR outlet has been suitably adapted based on its heartbeat).

### Geometric reconstruction of the left heart internal wall surfaces

In this section, we describe a novel framework to reconstruct the LH geometry and displacement. This is based on combining two different reconstruction techniques proposed so far for LV and for LA/AR, respectively. The entire procedure is represented in Figure 1, from step A to step C.

Regarding the LV reconstruction, we adopted the strategy described in [32]. Starting from the Short-Axis series of LV, we merged them with the Long-Axis acquisitions (2CH, 3CH and 4CH views) to obtain new enhanced time-dependent series of volumetric images with a uniform space resolution of 1 *mm* in all directions. From these enhanced images, we segmented and reconstructed the shape of the LV endocardium in all the 30 frames by using the algorithm proposed in [39] and implemented in the Medical Image Toolkit (MITK) open-source software (www.mitk.org), see step A in Figure 1.

After, we reconstructed the shape of AR and LA in all the 30 frames by using a cine-MRI multi-image based reconstruction algorithm, proposed in [37] for the right heart and implemented in the Vascular Modeling Toolkit (VMTK) (www.vmtk.org) [40, 41]. This is based on manually tracing the contours of AR and LA from the Rotated series of MV, the Long Axis series, and, when available, the Short Axis of AR. For each frame, a 3D point cloud was obtained, that was turned into a surface mesh of triangles, see step B in Figure 1.

Then, for each reconstructed frame, we merged the geometry of LV with the AR and LA ones in MeshMixer (https://www.meshmixer.com to obtain the final surface mesh of LH, see Figure 1, step C. Subsequently, we registered the displacement of each frame with respect to the end systolic configuration by exploiting the non-affine B-splines algorithm implemented in the Elastix open source library (http://elastix.isi.uu.nl) [42] used and validated in [35]. The output is the surface mesh of LH at the end systolic instant, where the displacement field 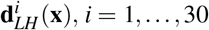, is defined for all the 30 frames. In Figure 1, step C, we reported the magnitude of 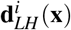 at three representative frames.

### Geometric reconstruction of the valves

In this section, we describe the geometric reconstruction of the valves. The entire procedure is reported in Figure 1, from step D to E. Regarding the MV reconstruction, starting from the Rotated series of MV, we reconstructed its shape in the fully closed (FC) and fully open (FO) configurations by using the method proposed in [38] (see step D in Figure 1), where the authors performed structural analysis, see also [43]. This method is based on tracing the valve leaflets in each plane to obtain a 3D point cloud that was after fitted with a B-Spline and then turned into a surface mesh of triangles in Matlab (www.mathworks.com), see [35] for details, where this MV reconstruction procedure has been, for the first time, applied to DIB-CFD simulations.

Concerning AV, the cine-MRI images at disposal did not allow a complete reconstruction of the patient-specific leaflets, thus we exploited a FO aortic valve template taken from the Zygote solid 3D heart model, a complete geometry reconstructed from CT scans representing an average healthy heart (https://www.zygote.com). We adapted it to our patients by exploiting the available structures that we were able to reconstruct. In particular, the FO configuration was geometrically deformed in order to match the annulus with that segmented from our cine-MRI images (from the AR Short Axis series for subject H and from the Rotated series of MV for patient R), see step F in Figure 1. The AV FC configuration was obtained for subject H by means of the same procedure used for FO, while for patient R it was obtained by exploiting algorithms based on the closest-point distances [41], see step F in Figure 1.

### Mathematical and numerical modeling

Blood was modeled as an incompressible, homogeneous, Newtonian fluid with density *ρ* = 1.06 10^3^ *kg/m*^3^ and dynamic viscosity µ = 3.5·10^*−*3^ *Pa·s*, described by the Navier-Stokes (NS) equations, see [44, 45]. To solve NS in moving boundaries we used the Arbitrary Lagrangian Eulerian (ALE) framework [46]. To treat the presence of the valves, we used the Resistive Immersed Implicit Surface (RIIS) method [47,48] and to evaluate the transition to turbulence, we employed the σ-LES method proposed for ventricular blood dynamics [49] and successfully used in different hemodynamic applications [50,51]. The entire method is described in a previous study [35]. In particular, the displacement of LH 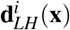 is used to compute the wall velocity to prescribe as boundary condition for the NS equations. Since 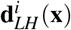 has been obtained only at the MRI acquisition times (*i* = 1, …, 30), we employed a spline interpolation to obtain **d**_*LH*_(**x**, *t*) for all *t* ∈ [0, *T*] where *T* is the duration of the heartbeat, equal to 0.9 *s* for subject H and 0.8 *s* for patient R, see Table 1. According to the ALE framework, at each time, the fluid domain Ω(*t*) is obtained by extending **d**_*LH*_(**x**, *t*) into Ω through the solution of a linear elastic problem [52].

The entire domain with its boundaries is displayed in Figure 2B. In particular, Σ_*LH*_ represents the internal wall surfaces of LH, Σ_*AR*_ and Σ_*PV*_, the final sections of the aortic root and pulmonary veins, respectively. In green, instead, we reported the surfaces of the aortic Γ_*AV*_ and mitral valve Γ_*MV*_.

Thus, the ALE NS equations in the known domain Ω(*t*) are solved to find the pressure *p* and the blood velocity **u**:

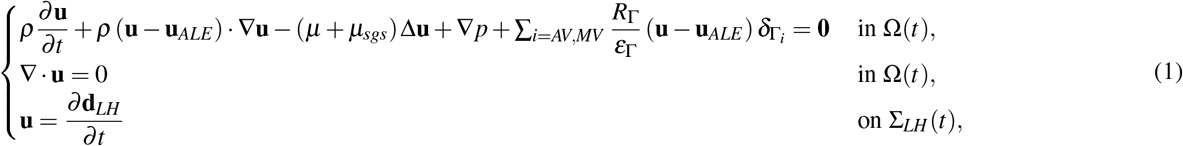

with a null initial condition in Ω(0). µ_*sgs*_ is the sub-grid viscosity of the σ-model [49], whereas 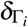 is a smoothed Dirac delta function representing a layer, with thickness 2ε_Γ_, around the surface of the valve Γ_*i*_, *i* = *AV, MV*, [31,48], and *R*_Γ_ is a resistance coefficient. In our numerical experiments, we set *R*_Γ_ = 10^5^ *kg/m·s* and ε_Γ_ = 0.75 *mm* [35, 53].

The valve dynamics has been modeled in an on-off modality, where the reconstructed leaflets opened and closed instantaneously according to the following rule [54]:

- if Δ*P >* 0 *→* valve opens,
- if *Q*_*AV*_ *<* 0 *→* AV closes, *→*
- if *Q*_*MV*_ *<* 0 & *t >* 0.77 *s→* MV closes^1^,

where Δ*P* is the difference between upstream and downstream pressures and *Q*_*AV*_ and *Q*_*MV*_ are the flow rates through AV and MV, respectively. Moreover, in order to guarantee a perfect adhesion between the valves and LH internal wall surfaces, we imposed that both valves move in accordance with the ALE movement of LH.

Regarding the remaining boundary conditions of system (1), we prescribed a Neumann condition in the normal direction by imposing on Σ_*PV*_ a constant pressure of 10 *mmHg* for H [55, 56] and a time dependent evolution for R [16], see Figure 2C, whereas a time dependent physiological trend representing the aortic pressure [16, 56] at Σ_*AR*_ for both cases, see Figure 2C. In the tangential direction, in order to avoid possible backflows instabilities, we prescribed a null velocity [57].

To numerically solve system (1), we used first-order Finite Elements together with fist order semi-implicit discretization in time [54]. The numerical scheme was stabilized by means of a Streamline Upwind Petrov-Galerkin/Pressure-Stabilizing

Petrov-Galerkin (SUPG/PSPG) scheme [58] implemented in the multiphysics high performance library *li f e*^*x*^ [59] (https://lifex.gitlab.io/) based on the deal.II core [60]. We run the simulations using 384 parallel processes on the GALILEO100 supercomputer (https://www.hpc.cineca.it/hardware/galileo100) at the CINECA high-performance computing center (Italy).

Tetrahedral meshes were generated in VMTK with an average mesh element size equal to 1.1 *mm* for H and 1.5 *mm* for R, with a local refinement of 0.3 *mm* close to valves, corresponding to about 1.8M degrees of freedom in both the cases (see Figure 2B for the mesh of R). The timestep Δ*t* was equal to 5 · 10^*−*4^ *s*. We performed a mesh convergence test ensuring that no significant differences may be found by using a finer mesh or a smaller timestep.

We simulated 13 heartbeats and we discarded the first one to remove the influence of the null initial condition.

### Quantities of interest

To describe and quantify the transition to turbulence, risk of hemolysis and thrombi formation in the two scenarios, we quantitatively analyzed and evaluated pressure and velocity, solutions of (1). We referred to them as ensemble quantities, calculated over 12 heartbeats. Moreover, we calculated also other quantities obtained by a post-processing of the ensemble velocity:

- wall shear stresses (WSS), that is a function of space and time representing the viscous forces, per unit of area, exerted by the blood on the walls [61]. In particular, we computed the time average wall shear stresses, TAWSS(**x**). High values of TAWSS may damage the LH endocardium and trigger possible remodeling processes [62];
- relative residence time (RRT), that is a function of space measuring the total time a fluid particle spends on the LH internal wall surfaces. High values of this quantity may be a marker of stagnant flow [63];
- E-wave propagation index (EPI), that is the ratio between the space covered by the blood jet developing at the MV orifice during the E-wave and the length of the ventricle at the end diastolic configuration. Values of EPI < 1 could indicate an incomplete apical washout, leading to possible left ventricle thrombus formation [64];
- standard deviation (SD) at each time and space of the blood velocity with respect to its ensemble value. This allowed us to quantify and localize the regions characterized by marked transition to turbulence [50, 51];
- Global Turbulent Kinetic Energy (GTKE, known also as Integrated Fluctuating Kinetic Energy) at each time, quantifying the velocity fluctuations by means of the fluid Reynolds stress tensor [30, 65];
- maximum tangential stress τ_*max*_ of the fluid Reynolds stress tensor [65], that is a function of space and time quantifying the fluctuating (turbulent) forces exerted among the fluid layers and thus possible damage caused to blood cells. Notice that values greater than 800 *Pa* are considered to create the conditions that promote hemolysis [66].

## Results

In Figure 3A, we reported the trend in time of the mean ensemble ventricular (LV), aortic (AR) and atrial (LA) pressures in the blue, red and magenta spheres, respectively. Notice that the maximum systolic drop Δ*P*_*AV*_ between LV and AR pressures was reached at the middle of systole for H and slightly later (about 60 *ms*) for R, with values of 22 *mmHg* and 14 *mmHg*, respectively. As expected, R featured a lower Δ*P*_*AV*_ with respect to H, due to the presence of regurgitation [67]. The closure of the aortic valve occurred at 0.30 *s* for H (i.e. at 33% of the heartbeat) and at 0.32 *s* for R (i.e. at 40% of the heartbeat). During the diastolic phase, the ventricle pressure remained almost constant in H, with a peak of pressure drop between LA and LV Δ*P*_*MV*_ during the E-wave equal to 3.6 *mmHg* and 12.5 *mmHg* for H and R, respectively. The larger peak of Δ*P*_*MV*_ featured by R with respect to H was in accordance with [68]. The closure of the mitral valve occurred at 0.9 *s* and 0.8 *s* for H and R, respectively. Regarding the atrial pressure, in accordance to the boundary conditions imposed at PVs, we observed in the healthy case a constant value of 10 *mmHg* during the whole heartbeat, whereas, in the regurgitant scenario the pressure increased up to 40 *mmHg*. In Figure 3B, we reported the ensemble flow rates evaluated through AV (blue plane) and MV (red plane) for the two scenarios. During the systolic phase, the AV flow rate reached a maximum of 617 *mL/s* for H and 530 *mL/s* for R. Notice that the peak of the AV flow rate was reached at the same instant of maximum systolic Δ*P*_*AV*_ in both the scenarios. Moreover, the flow rate through MV in R featured a peak of 574 *mL/s*. During diastole, in the trend of MV flow rate we recognized the E-wave (first minimum), the A-wave (second minimum) and the diastasis (middle stage of diastole). The E-wave featured a maximum absolute value of 650 *mL/s* and 1525 *mL/s* for H and R, respectively. The higher diastolic MV flow rate in R was in accordance with [69]. During diastasis, the MV flow rate decelerated until a slightly reversal at 0.63 *s* (i.e. 70% of the heartbeat) and 0.60 *s* (i.e. 75% of the heartbeat) for H and R, respectively. During the A-wave, the flow rate reached absolute values of 190 *mL/s* for H and 392 *mL/s* for R. After the A-wave, the second flow reversal through MV occurred at 0.77 *s* (i.e. 96% of the heartbeat) in the regurgitant scenario.

**Figure 3.**
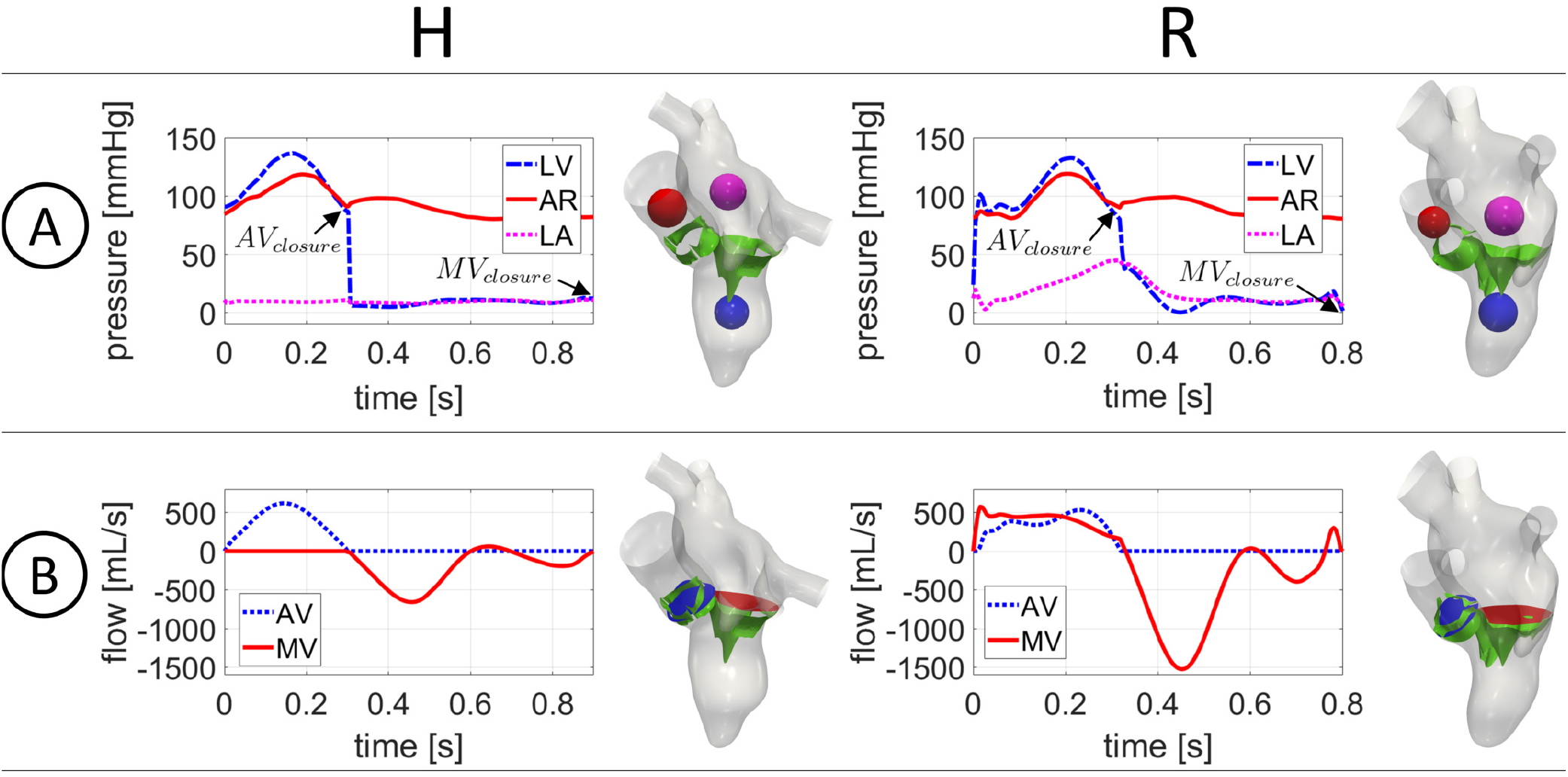
For each scenario: A: trend in time of the mean ensemble ventricle (LV), aortic (AR) and atrial (LA) pressures in the blue, red and magenta spheres, respectively; B: trend in time of the ensemble flow rates through AV (blue section) and MV (red section). *AV*_*closure*_: closure of aortic valve; *MV*_*closure*_: closure of mitral valve.

In Figure 4, we reported the magnitude of the ensemble velocity field on a slice along the 3CH axis at three representative time instants: the instant of peak systole (i.e. maximum AV flow rate) *t*_*PS*_, of peak E-wave *t*_*EW*_, and of peak A-wave *t*_*AW*_. In particular, *t*_*PS*_ = 0.15 *s* and 0.21 *s, t*_*EW*_ = 0.45 *s* and 0.465 *s, t*_*AW*_ = 0.8 *s* and 0.7 *s*, for H and R, respectively. At *t*_*PS*_ we displayed also a slice along the 2CH axis (see the black box), to better highlight that part of the ventricle flow went in the atrium resulting in the formation of a regurgitant jet developing along the anterior leaflet and along the LA walls, with a velocity peak of 5.5 *m/s* at the level of the MV orifice. The maximum velocity through the AV plane was equal to 2.1 *m/s* for H and 1.5 *m/s* for R. As expected, the peak of AV velocity was higher in the healthy scenario [16, 34]. Notice also the different velocity distributions in the atrium in the two scenarios: in H, no specific velocity patterns were observed, whereas in R the regurgitation promoted chaotic and irregular structure formations. At *t*_*EW*_, when the blood flow went from LA to LV, we obtained maximum velocity values through MV equal to 1.08 *m/s* and 1.73 *m/s* for H and R, respectively, highlighting the more elevated velocity in R, due to the higher MV flow rate, see Figure 3B. Furthermore, in both the scenarios we observed the formation of a ventricular vortex ring developing below the anterior leaflet. Moreover, during the E-wave we calculated the value of EPI in the ventricle, which was equal to 1 and 2 for H and R, respectively, highlighting the better ability of R to washout ventricular blood than H. At *t*_*AW*_, the second injection of fluid in the ventricle occurred. The velocity through MV were lower with respect to the ones observed at *t*_*EW*_ for both the scenarios. Furthermore, we noticed swirling structures in the ventricle, especially in correspondence of the middle-apex areas, due to vortices formed during the diastasis.

**Figure 4.**
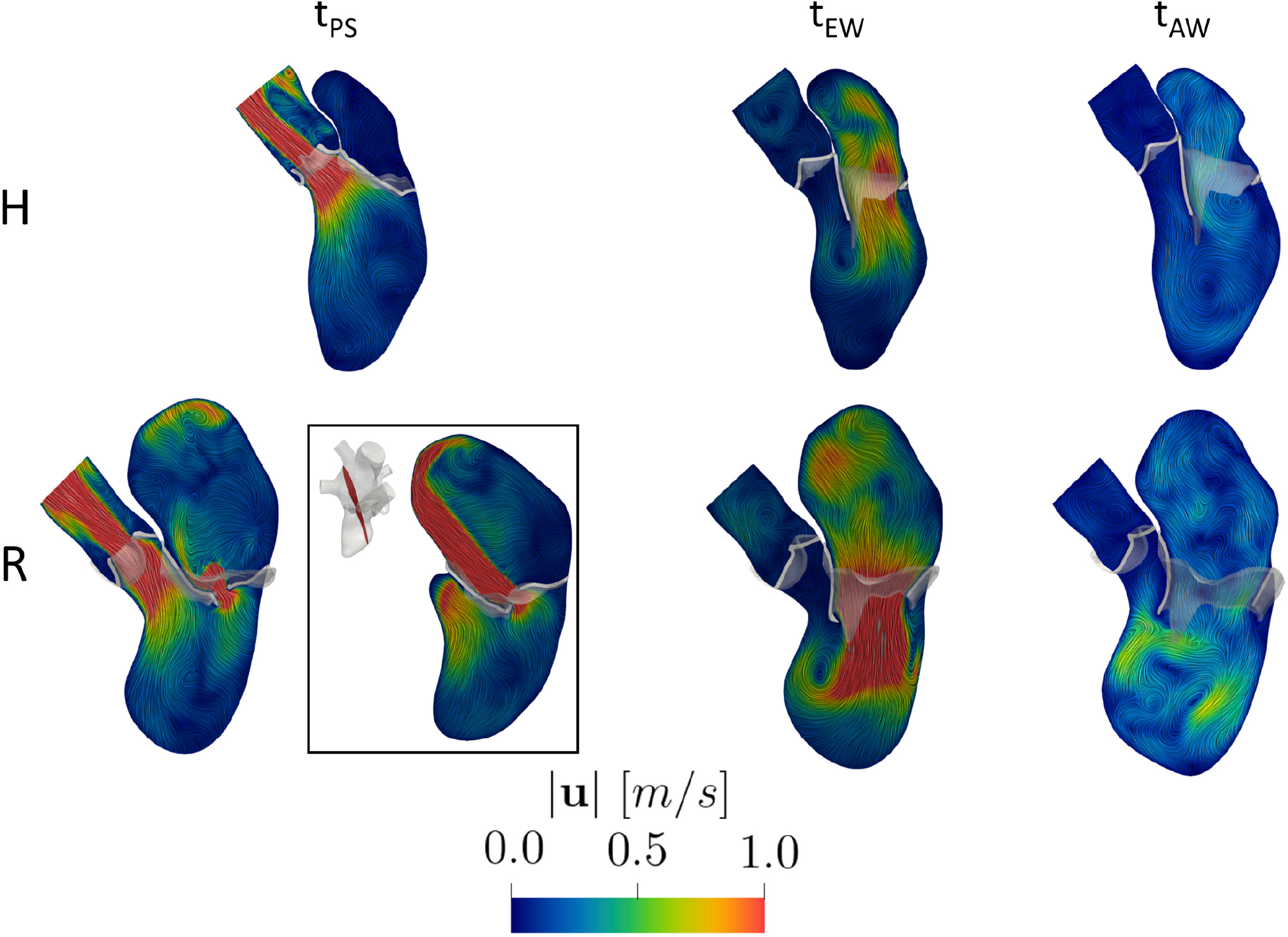
Magnitude of the ensemble velocity at three representative time instants over a slice along the 3CH axis in the two scenarios. At *t*_*PS*_, we also reported a slice along the 2CH axis (black box) for the regurgitant case.

In Figure 5, we reported the volume rendering of the ensemble velocity magnitude at *t*_*PS*_ (panel A), the spatial distribution of TAWSS (panel B) and RRT (panel C) for the two scenarios, in two different views. From this figure we observed that for R the regurgitant jet gave rise to high velocities, elevated values of TAWSS, and low values of RRT where the blood flow scratched and impinged against the atrial walls. Notice also that in the ventricle, TAWSS was slightly higher in R due to larger velocities occurred during the diastolic phase, see Figure 4. To quantify these differences, we reported in Table 2, the percentage of area with RRT greater than 5 *Pa*^*−*1^, computed over the total LV surface, the LV apex, the total LA surface, and left atrial appendage (LAA), in the two scenarios. Notice that such values were in any case larger for H than R. We pointed out that this threshold was chosen as representative value to discriminate high and low values of RRT. However, the analysis performed with other thresholds led to the same conclusions (percentage of area above the threshold larger in H).

**Table 2.** Values of the quantities of interest computed for each scenario. EPI: E-wave propagation index; Percentage of area with RRT greater than 5 *Pa*^*−*1^ evaluated in four different locations (LV, *LV*_*apex*_, LA, and LAA); *GTKE*: average in time of GTKE evaluated in LV and LA; Total exposure time of the blood to values of τ_*max*_ > 800 *Pa*.

| Scenario | $EPI$ [–] | $AREA\ RRT > 5\ Pa^{-1}$ [%] | | | | $\overline{GTKE}$ [mJ] | | $t\ \tau_{max} > 800\ Pa$ [ms] |
| --- | --- | --- | --- | --- | --- | --- | --- | --- |
| | | LV | $LV_{apex}$ | LA | LAA | LV | LA | |
| H | 1 | 68 | 83 | 48 | 100 | 0.20 | 0.12 | 0 |
| R | 2 | 40 | 53 | 33 | 100 | 4.80 | 7.30 | 220 |

**Figure 5.**
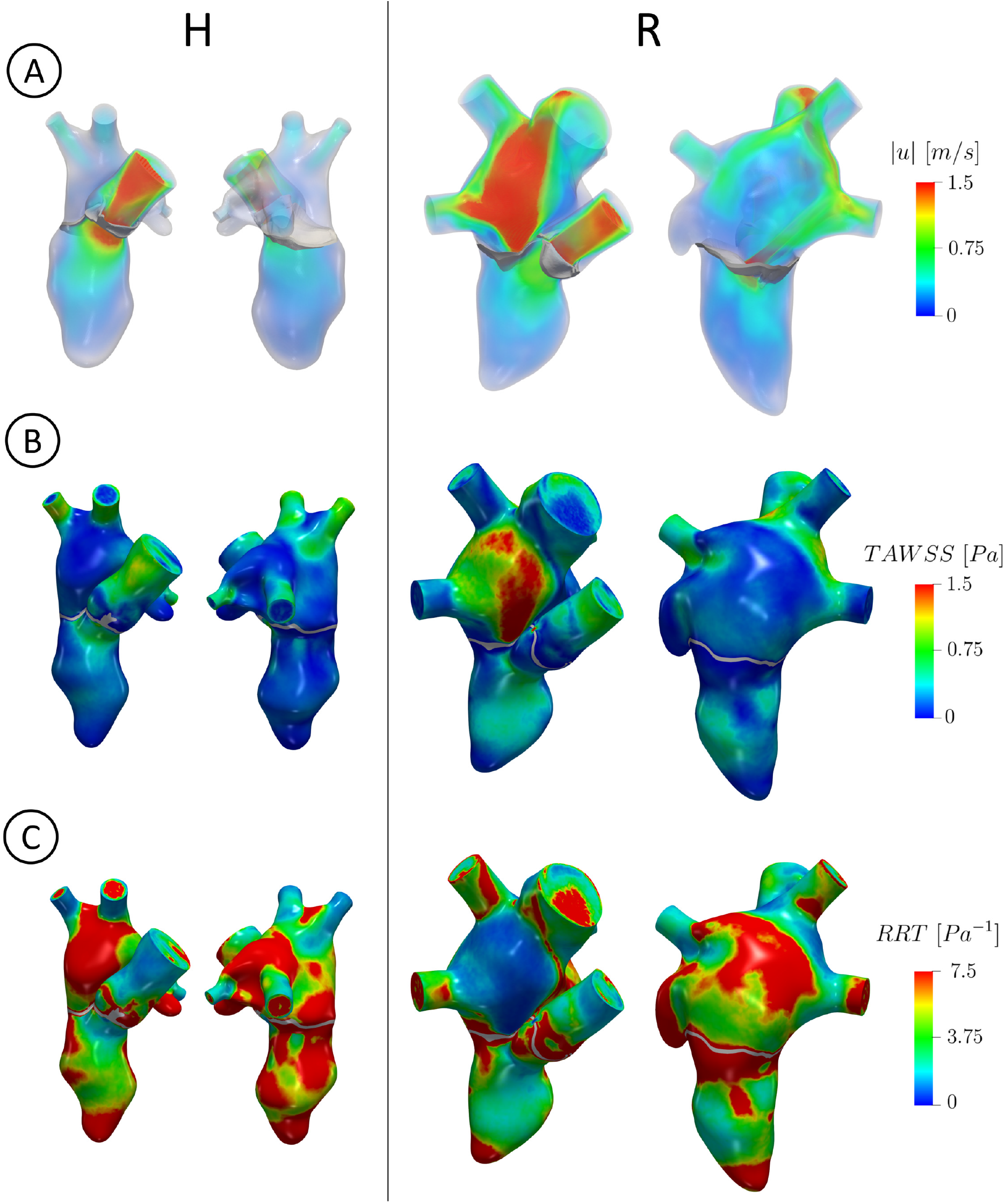
For each scenario in two different views: A: volume rendering of the ensemble velocity magnitude at *t*_*PS*_; B: spatial distribution of TAWSS, reported in the end systolic configuration; C: spatial distribution of RRT, reported in the end systolic configuration.

In Figure 6A, we reported, for each scenario, the evolution in time of GTKE evaluated in LV and LA. We observed that in R GTKE was much larger than in H in all the two chambers, with maximum values in LA at the end of systole due to the regurgitant jet and almost constant values in LV. With *t*_*GTKE*_ we referred to the time instant of maximum GTKE, equal to 0.29 *s* (i.e. at 32% of the heartbeat) and to 0.25 *s* (i.e. at 31% of the heartbeat) for H and R, respectively. In Table 2, we reported the average in time of GTKE, for both the scenarios confirming larger values for R. In Figure 6B, we displayed at *t*_*GTKE*_ and *t*_*EW*_ two slices along the 2CH (left) and 3CH (right) axes with the velocity SD. R featured in any case larger values of SD in all the left heart than H, confirming the greater predisposition of R to develop transition to turbulence. SD values in R at *t*_*GTKE*_ were comparable with the ensemble velocity ones, with a peak of 183 *cm/s* in correspondence of the regurgitant jet and of 39 *cm/s* in LV. Instead in H, the maximum SD value was equal to 17 *cm/s* in LA and to 19 *cm/s* in LV. Notice also a peak of 31 *cm/s* located in AR. At *t*_*EW*_, the fluctuations were mainly present in the center of LA with a peak of 55 *cm/s* in R. Instead in H, we noticed very low SD values with a peak in the LA center of 13 *cm/s*.

**Figure 6.**
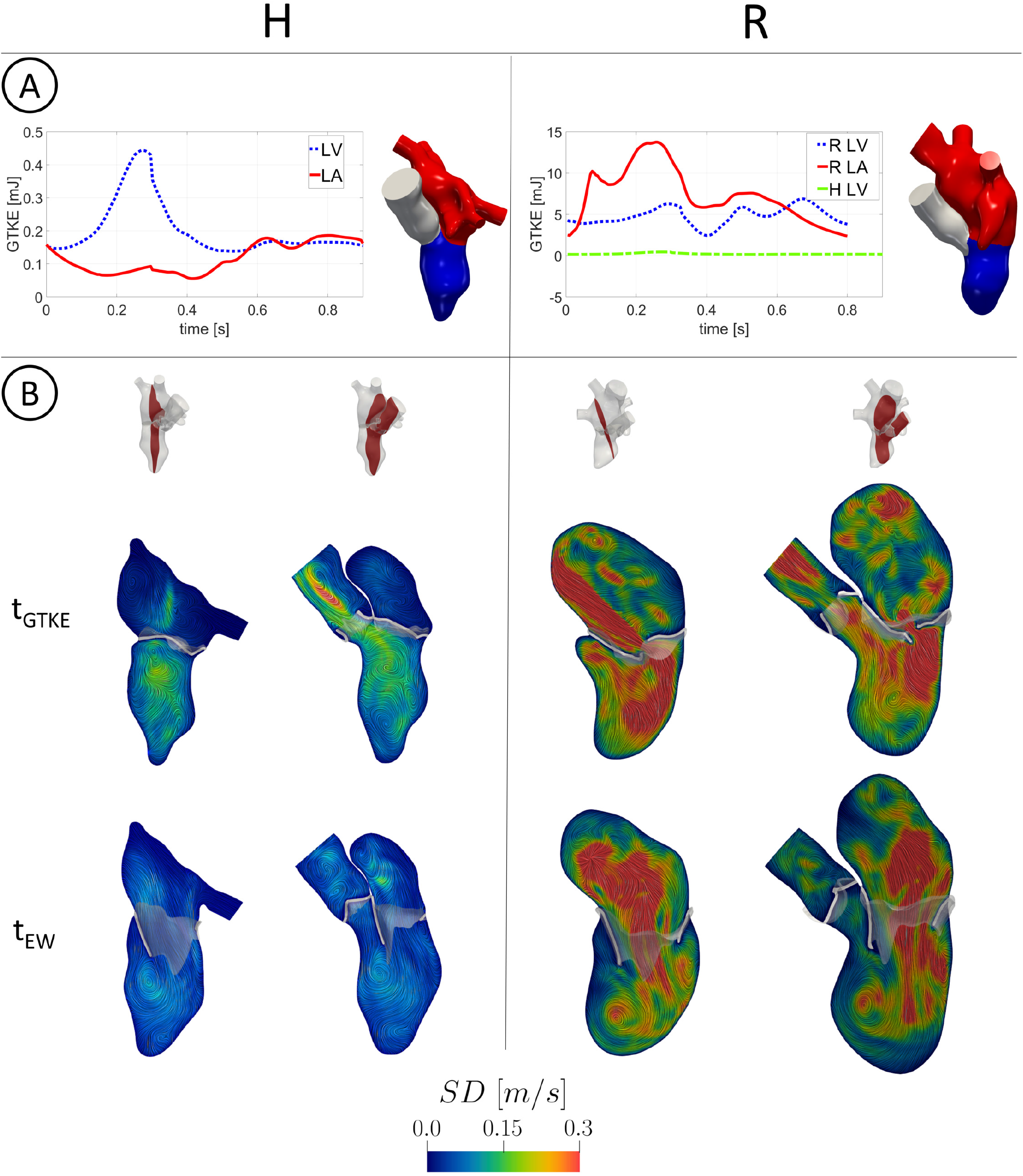
A: trend in time of GTKE evaluated in LV and LA. Notice the different scale used for H (left) and R (right). In the right figure, also the H LV case has been reported for a direct comparison; B: slices along the 2CH (left) and 3CH (right) axis with the velocity SD at *t*_*GTKE*_ and *t*_*EW*_.

In Figure 7A, we reported for the regurgitant scenario the volume rendering of τ_*max*_ at the instants where the volume of blood characterized by τ_*max*_ *>* 800 *Pa* featured its peaks, *t*_1_ = 0.07 *s* and *t*_2_ = 0.24 *s*, see Figure 7B. We observed values of τ_*max*_ greater than 800 *Pa* when the regurgitant jet impinged against the atrial walls (*t*_1_) and when a rapid deceleration of the regurgitant flow through MV occurred (*t*_2_), see also Figure 3B. From Figure 7B, we observed that during the diastolic phase the volume with elevated τ_*max*_ was always null. We also computed the total exposure time to values greater than 800 *Pa*, which was equal to 220 *ms* for R and 0 *ms* for H, suggesting the absence of hemolysis in the healthy scenario (see Table 2).

**Figure 7.**
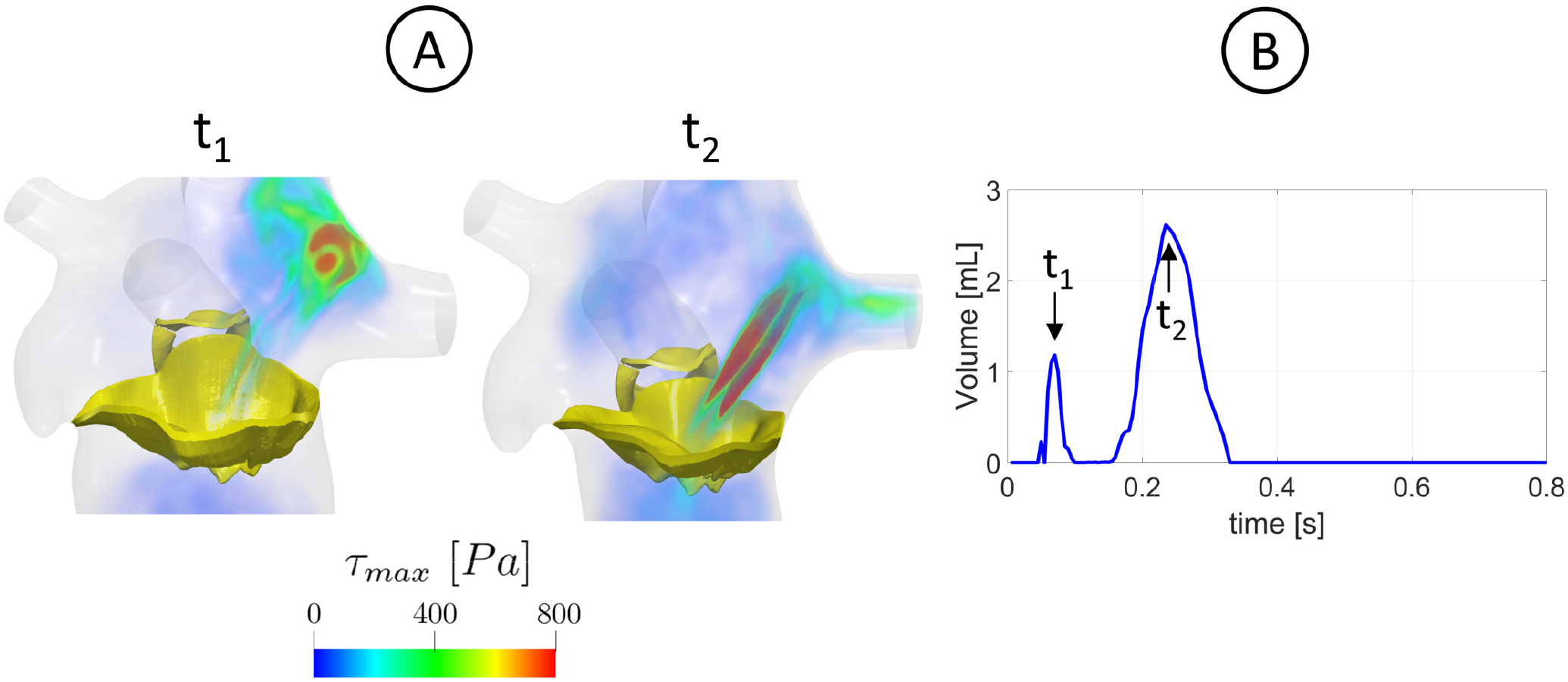
A: volume rendering at two different time instants *t*_1_ and *t*_2_ of the maximum tangential stress τ_*max*_ quantifying the possible damage caused to blood cells in the regurgitant scenario; B: time evolution of the volume of blood characterized by τ_*max*_ *>* 800 *Pa* and identification of *t*_1_ and *t*_2_ as the peak instants.

In Figure 2A, we reported, for the healthy scenario, the three locations of the acquisitions of ECD peak velocity measures. In particular, to assess the accuracy of the results of subject H reported in this work we compared the velocity field computed by the numerical simulation with such measures. In Table 3, we reported the corresponding values in the three locations. The maximum relative error Δ was found in P2 (5.9%), whereas in P1 and P3 the error was no larger than 4.0%.

**Table 3.** Comparison for the healthy case between the measurement of ECD and the values of the numerical simulation (SIM) in the three points of Figure 2A. Δ is the relative discrepancy.

| Scenario | P1 |  |  | P2 |  |  | P3 |  |  |
| --- | --- | --- | --- | --- | --- | --- | --- | --- | --- |
| | $ECD [m/s]$ | $SIM [m/s]$ | $\Delta [\%]$ | $ECD [m/s]$ | $SIM [m/s]$ | $\Delta [\%]$ | $ECD [m/s]$ | $SIM [m/s]$ | $\Delta [\%]$ |
| H | 1.25 | 1.20 | 4.0 | 0.85 | 0.80 | 5.9 | 0.62 | 0.60 | 3.2 |

In Figure 8, we reported a qualitative comparison of the regurgitant flow pattern obtained by the numerical simulation (bottom) with cine-MRI (top) at the three representative frames in R. In particular, in the cine-MRI views the regurgitant jet Validation of the results of subject H was detected by the areas characterized by a darker color, representing high blood flow velocities. Notice the good qualitative agreement between computations and images, highlighting that the regurgitant jet firstly developed along the anterior leaflet (Frames 1 and 2) and then also along the atrial walls assuming also a swirling structure (Frame 3).

**Figure 8.**
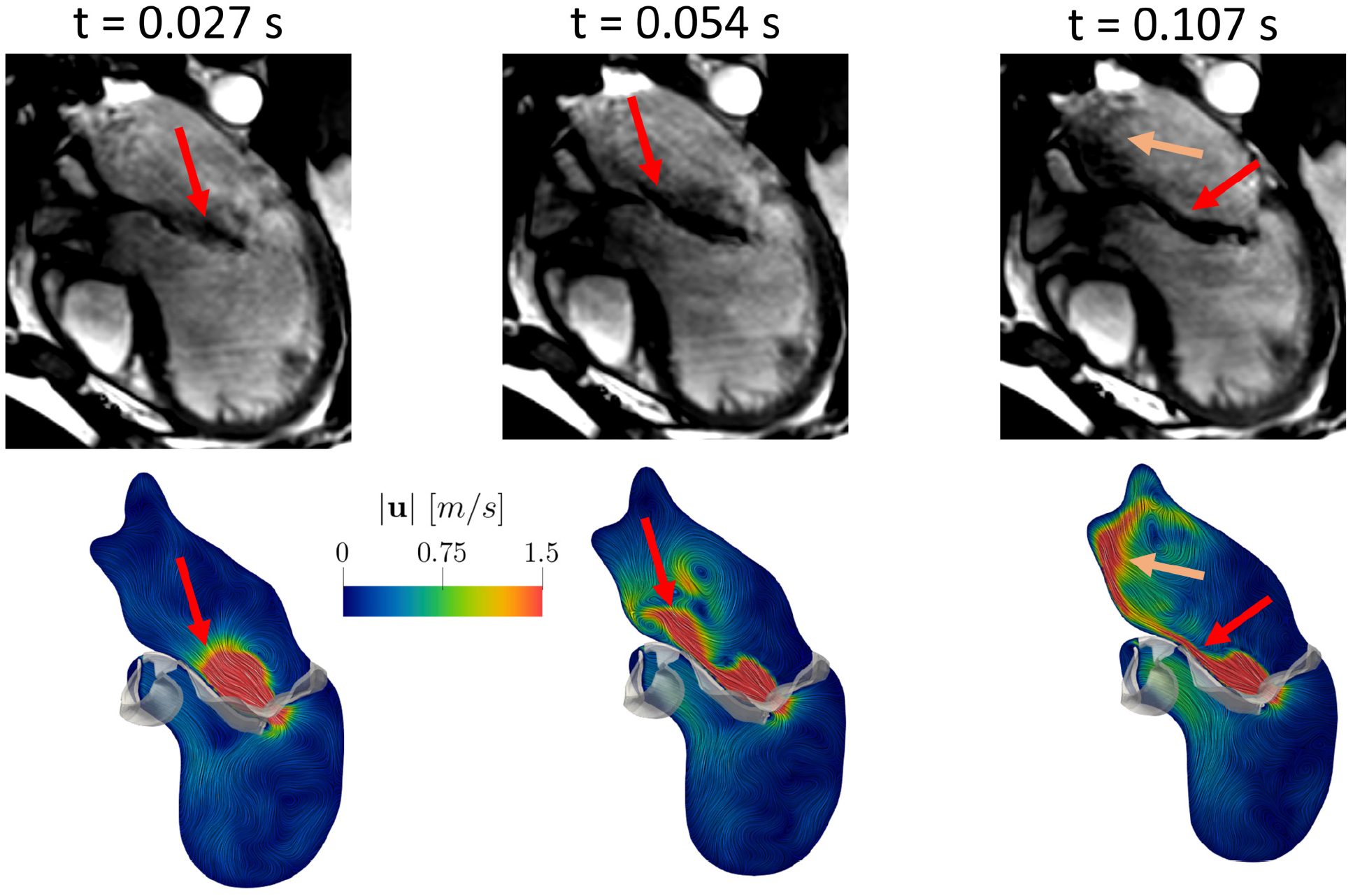
Comparison of the ensemble velocity pattern with Cine-MRI images at three representative frames in the regurgitant scenario. The arrows highlight the good agreement between computations and measures.

## Discussion

In this work we performed an image-based computational fluid dynamic study in the left heart of a healthy subject and of a patient with a severe MVR, with the aim of comparing the transition to turbulence, the risk of hemolysis, and the risk of thrombi formation. The motion of the complete LH (LV, LA, and AR internal wall surfaces, together with AV and MV) during the whole heartbeat was reconstructed and prescribed to DIB-CFD simulations from cine-MRI images.

At the best of our knowledge, this is the first work using all the patient-specific geometry motion of LV, LA, AR, AV and MV surfaces in a DIB-CFD simulation. In particular, for the healthy case the majority of the previous DIB-CFD studies did not include LA or AR or employed template geometries [25, 26, 31, 32, 34, 35]. In [19, 27, 30] the authors reconstructed all the patient-specific LH walls of a healthy subject, where however MV and AV in [19] and AV in [27, 30] were geometrically modeled as planes in their FC configuration and disappeared in the FO configuration. For the regurgitant case, no patientspecific LA and MV motions could be found in previous studies.

In our work, we included both the valves and the entire reconstruction of the LH internal wall surfaces and valves was performed by combining different acquisitions of cine-MRI images at disposal: the LV Long Axis, the LV Short Axis, the AR Short Axis (when available) and the MV Rotated series. The latter stand out from the other series because they are based on a radial sampling (while the other series on a Cartesian sampling) which consist in 2D Long Axis series of 18 evenly rotated planes (one every 10 degree) around the axis passing through the MV center and aligned with the LV apex, proposed for the first time in [38] for a pure structural analysis. Such images represented advanced acquisitions not acquired in the clinical routine, see Figure 1, which allowed us to obtain detailed 3D geometry motion of LA, AR, MV and AV, in addition to ventricle motion for which the LV Short and Long Axis series are sufficient.

Owing to this specific and detailed imaging, we were able to quantitatively describe physiological and pathological/MVR mechanisms such as prevention from thrombi, transition to turbulence, and hemolysis formation. In particular, for the first time at the best of our knowledge, we investigated by means of a computational study how the regurgitant jet could promote the washing out of regions with stagnant flow in LA, reducing the risk of thrombosis with respect to a healthy scenario. Our results highlighted that the regurgitant jet scratched against the atrial walls, see Figure 5A, resulting in elevated values of TAWSS, see Figure 5B. This led to low values of RRT (Figure 5C) indicating that there is little stagnation in those regions and thus promoting the washing out of blood. This was also confirmed by the values reported in Table 2. Notice however that RRT in LAA was elevated in both the subjects, confirming that, even in MVR, LAA could be one of most common site for cardiac thrombus formtaion [70]. Moreover, we investigated the mechanism of washing out occurring in the ventricle during the E-wave. In particular, in the regurgitant scenario there was a significant LV apical washout due to high blood flow coming from MV, see Figs. 3B and 4 as confirmed by the value of EPI and of low RRT areas, see Table 2. The high velocities through MV allowed the blood to reach the ventricle apex rapidly and with high energy before diastasis, allowing to remove possible areas of stagnant flow [64]. Thus, MVR promoted a more relevant washout, with respect to the physiological case, of the blood flow in LA during the systolic phase and in LV during the E-wave, leading to regions more protected from the possible formation of clots. Such results are in agreement with clinical studies [4, 71, 72].

Second, we investigated the transition to turbulence in the MVR case. So far, at the best of our knowledge, the turbulent effects were investigated only in physiological LH [27, 30]. By employing a LES model, we showed that the regurgitant scenario featured high velocity fluctuations among the heartbeats, resulting in elevated velocity standard deviation, see Figure 6 and the values of GTKE reported in Table 2. In particular, the maximum fluctuations in LA occurred for R at late systole, as also reported in [2], with a GTKE peak value equal to 14 *mJ*, which falls in the range (13, 37) *mJ* found in [2]. Moreover, the ratio of the average GTKE in LA between R and H was equal to 61, see Table 2, confirming that the presence of the regurgitant jet promoted more turbulence in R. We noticed also elevated transition to turbulence for R in LV, especially during the rapid deceleration of the systolic blood flow (Figure 3B), during the E-wave due to the formation of a ventricular vortex ring and to the impingement of the diastolic jet against the LV apex (Figure 4), and during the A-Wave due to the mixing of blood after diastasis (Figure 4). The ratio of the average GTKE in LV between R and H was equal to 24, highlighting how the regurgitation promoted a great amount of turbulence also in LV. We point out that the transition to turbulence in LV was more pronounced for R because of the larger heart dimension resulting in higher ventricular blood flow. Since the BSA of the two subjects was very similar (see Table 1), we argue that the larger LH dimensions of patient R were due uniquely to MVR, which is known to be correlated with progressive remodeling and dilation [73]. In this respect, we noticed that the ratio between R and H LV diameters at the end diastolic configuration was equal to 1.25, a value in agreement with medical studies reporting that severe MVR dilated the left ventricle diameter by a factor of 25-30% [74, 75].

Third, we observed that the definition of τ_*max*_ (see Section “Quantities of interest”) implies that large transition to turbulence (and thus large values of τ_*max*_) promotes hemolysis. In particular, we found high values of τ_*max*_ (greater than 800 *Pa*, identified as a risk threshold [66]) in correspondence of the regurgitant jet, see Figure 7A. We highlighted two different mechanisms (Figure 7B) that provoke large τ_*max*_: at *t*_1_ the fragmentation of the regurgitant jet against the atrial walls; at *t*_2_ the rapid deceleration of the blood flow through MV, see also Figure 3B. These two different mechanisms were also described by clinical studies [3, 76]. Furthermore, according to [66], regions exposed to value of τ_*max*_ larger than 800 *Pa* for more than 1 *ms* could experience the conditions of promoting hemolysis. In our R case, we found a value of 220 *ms*, see Figure 7B, and Table 2.

The blood velocity results found in this work for the healthy case have been validated against ECD measures acquired in the subject in three different locations. In particular, although far to establish a complete validation, they are very promising since they highlighted a good accuracy (with a maximum error of 6%, see Table 3) between the maximum values. Furthermore, we reported a qualitative comparison of the regurgitant flow pattern obtained by the numerical simulation with the cine-MRI, see Figure 8. We noticed a good agreement between the cine-MRI images and the numerical simulations. In particular, the direction of the regurgitant jet in the numerical simulation was also in accordance with the Carpentier’s functional classification for which, in case of MVR due to a prolapse, the jet is directed away from the pathological leaflet, in our case the posterior one [77, 78]. To support the other results and the physio-pathological implications, we compared our findings with the literature. In particular, in the healthy scenario the value of the maximum pressure drop Δ*P*_*MV*_ = 3.6 *mmHg* computed across MV falls down in the same range ≃ (2.0, 4.0) *mmHg* found in [79]. In the regurgitant scenario, we compared our velocity findings with measures in the case of a severe MVR. We found a value of the maximum velocity through MV at systole equal to 5.52 *m/s*, in accordance with the range (5.00, 6.00) *m/s* [2], and during the E-Wave equal to 1.73 *m/s*, in accordance with the range (1.18, 1.77) *m/s* [80]. Notice that the E-Wave peak velocity value is used in clinics to assess the severity of MVR. Notice also that the MV flow reversals detected at diastasis (*t ≃*0.59 *s*) and before the end of the heartbeat (*t ≃* 0.77 *s*, see Figure 3B) were also found in other computational studies [16, 19, 81].

## Limitations

Some limitations characterized this work:

1. We considered only two subjects. This was a consequence of the fact that we used advanced (not daily available) images series elaborated in order to perform highly accurate DIB-CFD simulations. This allowed us to extract interesting general physio-pathological findings of the healthy and regurgitant scenarios. As a consequence, our reconstruction technique is not suitable to study a wide range of specific subjects, for which standard images are enough;
2. We did not model the valve dynamics, instead we considered a instantaneous valve opening and closure. However, the opening and closing valve time duration is less than 5% of the heartbeat [82]. Thus, the opening and closure of the valves can be considered as instantaneous as a first approximation. The degree to which a patient-specific valve dynamics may affect the velocity field and the transition to turbulence is currently under study;
3. We did not include the subvalvular apparatus (papillary muscles + chordae tendineae) in the LV geometry. This is a common choice, adopted also in [26, 28, 29, 31, 32] due to the difficulty to reconstruct the papillary muscles and chordae tendineae from MRI images. However, their influence on the quantities of interest, in particular on transition to turbulence, should be relevant and it will be the subject of future studies;
4. We did not considered the MRI beat-to-beat variations in terms of acquired displacements and heart rate. Indeed, the average of MRI acquisitions along more heartbeats should provide more reliable input data for DIB-CFD. However, this may result in prolonged exposure time during the acquisitions of cine-MRI, that is not advisable for patients with heart pathologies.

## Acknowledgements

CV is partially supported by the Italian project MIUR PRIN17 2017AXL54F “Modeling the heart across the scales: from cardiac cells to the whole organ”. Moreover, the authors acknowledge the CINECA award under the ISCRA B initiative, for the availability of high performance computing resources and support (ISCRA grant *IsB*25_*MathBeat*, 2022-2023).

## Author contributions statement

Acquisition of the clinical data: GP, CM,VDN Methodology: LB, FR, CV

Image Reconstruction and numerical simulations: LB Conceptualization: LB, VG, VDN, GP, GBL, CV

Formal analysis and investigation: LB

Interpretation of the results: LB, CV

Writing - Original draft preparation: LB

Writing - Review and editing: CV, VG, FR, VDN, CM, GP, GBL

Supervision: CV, GBL

## Footnotes

1 ^1^The condition *t >* 0.77 *s* prevents the closure of MV during the flow reversals occurring before the end of the heartbeat, e.g at the diastasis, see Figure 3B in the Results section

